# Intracranial neurophysiological correlates of rumination

**DOI:** 10.1101/2024.05.08.593187

**Authors:** Xiao Chen, Zhen Fan, Dong Chen, Liang Wang, Liang Chen, Chao-Gan Yan

**Affiliations:** CAS Key Laboratory of Behavioral Science, Institute of Psychology, Beijing 100101, China; Temerty Centre for Therapeutic Brain Intervention, Campbell Family Research Institute, Centre for Addiction and Mental Health, Toronto M6J1H4, Ontario, Canada; Department of Psychology, University of Chinese Academy of Sciences, Beijing 100049, China; Magnetic Resonance Imaging Research Center, Institute of Psychology, Chinese Academy of Sciences, Beijing 100101, China; International Big-Data Center for Depression Research, Chinese Academy of Sciences, Beijing 100101, China; Department of Neurosurgery, Huashan Hospital, Shanghai Medical College, Fudan University, Shanghai 200040, China; National Centre for Neurological Disorders, Shanghai 200040, China; Shanghai Key Laboratory of Brain Function and Restoration and Neural Regeneration, Shanghai 200040, China; Neurosurgical Institute of Fudan University, Shanghai 200040, China; Shanghai Clinical Medical Centre of Neurosurgery, Shanghai, China, Shanghai 200040, China; CAS Key Laboratory of Mental Health, Institute of Psychology, Beijing 100101, China

**Keywords:** rumination, iEEG, precuneus, hippocampus, oscillation power

## Abstract

Rumination is a transdiagnostic psychological process that plays a prominent role in many common psychiatric disorders, albeit its neurophysiological basis remains elusive. Existing neuroimaging studies have highlighted the precuneus and hippocampus as two essential brain regions in rumination’s neural underpinnings. Here, we examined the intracranial electroencephalogram (iEEG) recordings from 21 patients with epilepsy during a naturalistic, continuous, active rumination state and measured the slow frequency (1-8 Hz) and high gamma (70-150 Hz) band oscillation powers. We observed enhanced slow frequency power in the precuneus and reduced high gamma power in the hippocampus during the rumination condition compared to the control condition. The hippocampal high gamma power reduction was associated with the self-reported reflection tendency. Our findings provided the first empirical evidence of the neurophysiological underpinnings of rumination and implicated a precuneus-hippocampus coupling across neural oscillation bands during an active rumination state.

## Introduction

Rumination is uncontrollable, self-reflective, and repetitive thinking about the distress and its possible causes and consequences (*1*). Even though it could be viewed as a transdiagnostic behavioral element in a number of common psychiatric disorders, most previous studies highlighted its close association with major depressive disorder (MDD) and its pivotal role in MDD’s psychopathology (*2, 3*). To date, rumination’s underlying neurophysiological basis remains elusive, and the existing evidence on its response to the current first-line antidepressant therapy is mixed (*4-7*). Accordingly, a better understanding of rumination’s neural basis may pave the way for the novel rumination-focused neuromodulation treatment of MDD.

Existing evidence from studies using functional magnetic resonance imaging (fMRI) has shown that brain regions of the default mode network (DMN) are involved in active rumination (*8-13*). Previous resting-state fMRI (rs-fMRI) studies have also revealed an association between self-reported ruminative tendency and the spontaneous functional coupling among DMN and other brain regions (*14-19*). Due to its self-enhanced nature and high self-constraint level, it is plausible to induce participants into a continuous, active “rumination state” and characterize its network underpinnings (*20*). Using this paradigm and fMRI, our previous studies and other research have found an altered functional connectivity pattern regarding the precuneus, one of DMN’s most critical nodes (*21-24*). One subcortical region that has been demonstrated to be closely associated with rumination is the hippocampus. The altered hippocampal activity was revealed when participants engaged in an active rumination (*12*). The self-reported ruminative tendency was related to brain activity in the hippocampus during an emotional processing or executive control task (*25, 26*). The functional coupling between the hippocampus and DMN has been shown to be associated with a ruminative tendency (*27*).

While existing functional brain imaging studies have highlighted the prominent role of the precuneus as one of the critical nodes of DMN and the hippocampus in the neural mechanisms underlying rumination, its neurophysiological basis remains largely unknown. The intracranial electroencephalogram (iEEG) recording provides both precise anatomical location and exceedingly high signal-to-noise ratio and may help delineate the electrophysiological basis of the human functional brain imaging findings. Here, leveraging the iEEG recordings from a group of patients with epilepsy, we intended to explore the electrophysiological features of the precuneus and hippocampus during an active rumination state. We used high gamma (70-150 Hz) and low frequency (1-8 Hz) as the surrogate measures of the local neurophysiological activities. High gamma power is reliably correlated to the BOLD signal (*28-31*), and the low-frequency local field potential (LFP) is involved with higher cognitive processes such as working memory and cognitive control (*32-35*). We hypothesized that these two regions would show different electrophysiological activity patterns during rumination compared to the control condition.

## Methods

Twenty-one patients with intractable epilepsy participated in this study. They had been surgically implanted with stereotactic electrodes as part of their pre-surgical assessment of seizure focus. Only right-handed patients and those who possess normal or corrected-to-normal vision were included in the current study. All electrode placements were decided solely based on clinical evaluation for surgery. Participants provided written consent to participate in the research, and the research protocol was approved by the Institutional Review Board of the Institute of Psychology, Chinese Academy of Sciences, and Huashan Hospital.

### Task description

Participants were asked to finish a rumination state task. Prior to the recording, participants were briefed on the purposes and requirements of the task. They also provided four groups of keywords. Each group denotes one adverse or pleasant autobiographical memory event that happened to them before. These keywords will be presented to them and serve as a reminder. This rumination task consists of five conditions. (1) Resting state: A white fixation (“+”) was presented on a black background, and the participants were asked to stay awake and not to think anything particular; (2) Sad memory: the keywords provided by the participants were shown on the screen and the participants were prompted to recall every detail of the corresponding event as they were re-experiencing it; (3) Rumination: prompts (e.g., “Think: Why can’t I handle things better in these events I just remembered?”) that are modified from the rumination inducement task (*11*) and the ruminative response scale (*36*) were presented and the participants were asked to reflect on themselves accordingly; (4) Distraction: participants were asked to imagine an objective scene as vividly as possible according to the prompts (e.g., “Think: A train stopped at a station”). (5) Happy memory: similar to the sad memory condition, participants were prompted to recall the corresponding autobiographical event in detail. This condition came right after the resting state or at last to counter-balance with the sad memory condition. The sad memory, happy memory, rumination, and distraction conditions consist of four sets of prompts. Each prompt lasted for 2 minutes. No inter-stimuli interval existed between two consecutive prompts, forming a continuous 8-minute state. The order between rumination and distraction was counterbalanced among all participants. Before the resting state and after all conditions, participants were asked to use a number to denote their emotional levels (1: very unhappy; 9: very happy). The content and form of the thoughts (e.g., negative vs. positive) in each condition were measured with a short 9-point questionnaire (1: not at all; 9: almost all) following our prior work (*24*).

### Self-report measures

A 22-item ruminative response scale (RRS) was administered to characterize participants’ trait rumination (*36*). This questionnaire contains two sub-scales: brooding, assessing a passive comparison to others or unrealistic standards; reflection, assessing a neutral self-focus (*37*). RRS has a good reliability (Cronbach’s alpha = 0.88 - 0.92) and validity (*38, 39*).

### Behavior Analysis

Participants’ ratings were analyzed using R (version 4.3.2) (*40*) with rstatix (*41*). A one-way ANOVA was performed to examine the Condition effect (fixed effect, before and after the resting state, sad memory, rumination, and distraction) on the self-reported emotion levels and the thinking content/forms. We performed post hoc pairwise comparisons among conditions using paired t-tests with the Bonferroni correction.

Recording methods

Recordings were performed at the Huashan Hospital using a standard clinical system (EEG-1200C, Nihon Kohden, Irvine, CA) with a 2000 Hz sampling rate. The implantation of electrodes was performed using 0.8 mm diameter electrode shafts consisting of 8-16 contacts. The contacts were 2 mm long and spaced with a 3.5 mm center-to-center distance. The contacts were labeled with consecutive numbers, from one (the deepest) to the most superficial. No seizure was observed 1 h before or after the tests in all patients.

We performed electrode localization with FieldTrip version 20221210 (*42*) run in the MATLAB R2023b (The MathWorks Inc., Natick, MA, US) platform. Generally speaking, the pre-implantation magnetic resonance imaging (MRI) images were first processed with Freesurfer’s recon -all (*43*) functionality. Participants’ post-implantation computed tomography (CT) images were then co-registered onto their pre-implantation MRI images. We visually identified electrode locations in each participant’s native space. Individual participants’ MRI images were then registered to the standard MNI coordinate space using a volume-based registration technique based on SPM 12 (*44*). We applied the resulting deformation parameters to the original electrode locations to obtain the locations in the MNI space. The standardized spatial positions of the electrodes were visually inspected to ensure the accuracy of the registration.

We defined regions of interest (ROIs: precuneus and hippocampus) according to the automated anatomical labeling (AAL) atlas (*45*). The anatomical labels of each electrode were assigned according to their locations in the MNI space.

### Preprocessing of the iEEG data

We first down-sampled the original iEEG signals to 1000 Hz and then band-pass filtered from 0.15 to 200 Hz using a two-pass Butterworth filter (ft preprocessing.m in the FieldTrip toolbox (*46*)). The raw data were notch-filtered at 50 Hz, and the harmonics were 50 Hz. A bipolar montage was applied to all the depth electrodes by taking bipolar derivations of consecutive channels.

We visually inspected the raw iEEG data and their corresponding power spectrum to rule out channels containing signal artifacts or inter-ictal epileptiform activity. Then, an automatic removal procedure was applied to the down-sampled raw iEEG data. The iEEG data’s envelope was obtained using a Hilbert transformation. The time points with epileptic discharges were identified according to the following criteria: (1) The envelope of the unfiltered signal exceeded the baseline by four standard deviations, where the baseline represents the signal’s overall mean value. (2) The filtered signal’s envelope, bandpass-filtered between 25 and 80 Hz, exceeded the baseline by five standard deviations, in line with prior research (*47*). Furthermore, channels with more than 50% of time points identified as noises were excluded from further analysis. The timing of all epileptiform activities and artifacts were stored for the following analysis.

### Time-frequency analysis

A power spectrum was obtained by performing five-cycle Morlet wavelets at 76 logarithmically spaced frequencies ranging from 0 to 150 Hz. This time-frequency transformation was performed using ft freqanalysis.m in the FieldTrip toolbox. One second before and after the time points identified as noises in the abovementioned automatic removal procedure were excluded. The power values were then *Z*-scored according to the average power at the channel level over all the interested conditions (rumination and distraction). Finally, the mean powers of 2 frequency bands (low frequency, 1-8 Hz and high gamma, 70-150Hz) were obtained by averaging all *Z*-standardized power value time series across all corresponding frequencies. Here, we combined delta (1-4 Hz) and theta (4-8 Hz) band frequencies as “low frequency” in light of the evidence implicating a common correspondence of these two bands to animal studies (*48, 49*). The mean power values of all channels in one specific ROI (precuneus and hippocampus) from one participant were averaged to get the subject-level mean power value for each frequency band. A paired *t*-test was performed on the subject-level mean power values. Given the well-established role of the hippocampus in autobiographical memory (*50*), we further compared the oscillation band power in the hippocampus between two autobiographical memory conditions (sad and happy) and the resting state, respectively.

### Association between trait rumination and neurophysiological features

Pearson’s correlation was used to explore the relationship between self-report rumination and the relative change of band powers between rumination and distraction conditions. Multiple comparisons were corrected with Bonferroni correction (2 ROIs × 2 bands). Four participants were excluded due to the missing values of self-reported questionnaires.

### Sensitivity analysis

We further compared the low-frequency band power and the high-gamma band power between rumination and distraction states in the middle occipital gyrus to test the sensitivity of our main findings. We performed paired *t*-tests to compare the mean power between each block of the rumination condition and the entire distraction condition to test the reliability of our main findings across all rumination prompts.

## Results

### Behavior data

Twenty-one patients (10 females; mean age = 28.67 years; age range: 17 to 60) participated in the current study (Table 1). The behavioral results showed that participants generally followed the task prompts. A one-way ANOVA revealed a significant condition effect (*F*(4, 80) = 13.89, *p* < .001, ***η***^2^ = .33). Post hoc comparisons showed that participants’ emotional ratings after the rumination condition were significantly lower than those after the distraction condition (*t*(20) = -4.07, *p*_*adj*_ = 0.01)/the resting state (*t*(20) = -3.32, *p*_*adj*_ = 0.03). Participants’ emotional levels after the sad memory condition were significantly lower than those before (*t*(20) = -5.46, *p*_*adj*_ < .001) and after the resting state (*t*(20) = -5.90, *p*_*adj*_ < .001)/the distraction state (*t*(20) = -4.92, *p*_*adj*_ < .001). Significant differences regarding the thinking contents were revealed among the rumination, distraction, and resting states except for the “future” and “others” dimensions (Figure 2ti and Table S2). Participants’ thinking contents in the rumination condition were more about the past and themselves, more negative, in the form of speech, and made them sadder compared to the distraction condition. On the contrary, participants thought less positively, less about the imagery, and less happily during rumination compared to distraction. tiompared to the resting state, participants thought more about the past, more negatively, less happily, and more sadly in the rumination condition.

**Table 1.** Demographic and clinical information of participants.

| **ID** | **Sex** | **Age**<br>**(year)** | **Number of channels**<br>**in hippocampus** | **Number of channels**<br>**in precuneus** |
| --- | --- | --- | --- | --- |
| Sub001 | Female | 23 | 11 | 0 |
| Sub002 | Female | 23 | 6 | 0 |
| Sub003 | Male | 32 | 13 | 0 |
| Sub004 | Male | 23 | 4 | 0 |
| Sub005 | Male | 26 | 3 | 6 |
| Sub006 | Male | 23 | 9 | 2 |
| Sub007 | Male | 29 | 17 | 0 |
| Sub008 | Male | 24 | 0 | 2 |
| Sub009 | Female | 25 | 4 | 0 |
| Sub010 | Male | 32 | 7 | 0 |
| Sub011 | Female | 27 | 11 | 2 |
| Sub012 | Female | 22 | 10 | 1 |
| Sub013 | Male | 32 | 12 | 0 |
| Sub014 | Female | 35 | 14 | 0 |
| Sub015 | Female | 27 | 13 | 3 |
| Sub016 | Female | 24 | 11 | 8 |
| Sub017 | Female | 60 | 14 | 0 |
| Sub018 | Female | 38 | 8 | 0 |
| Sub019 | Male | 17 | 0 | 12 |
| Sub020 | Male | 30 | 8 | 0 |
| Sub021 | Male | 30 | 11 | 0 |

**Figure 1.**
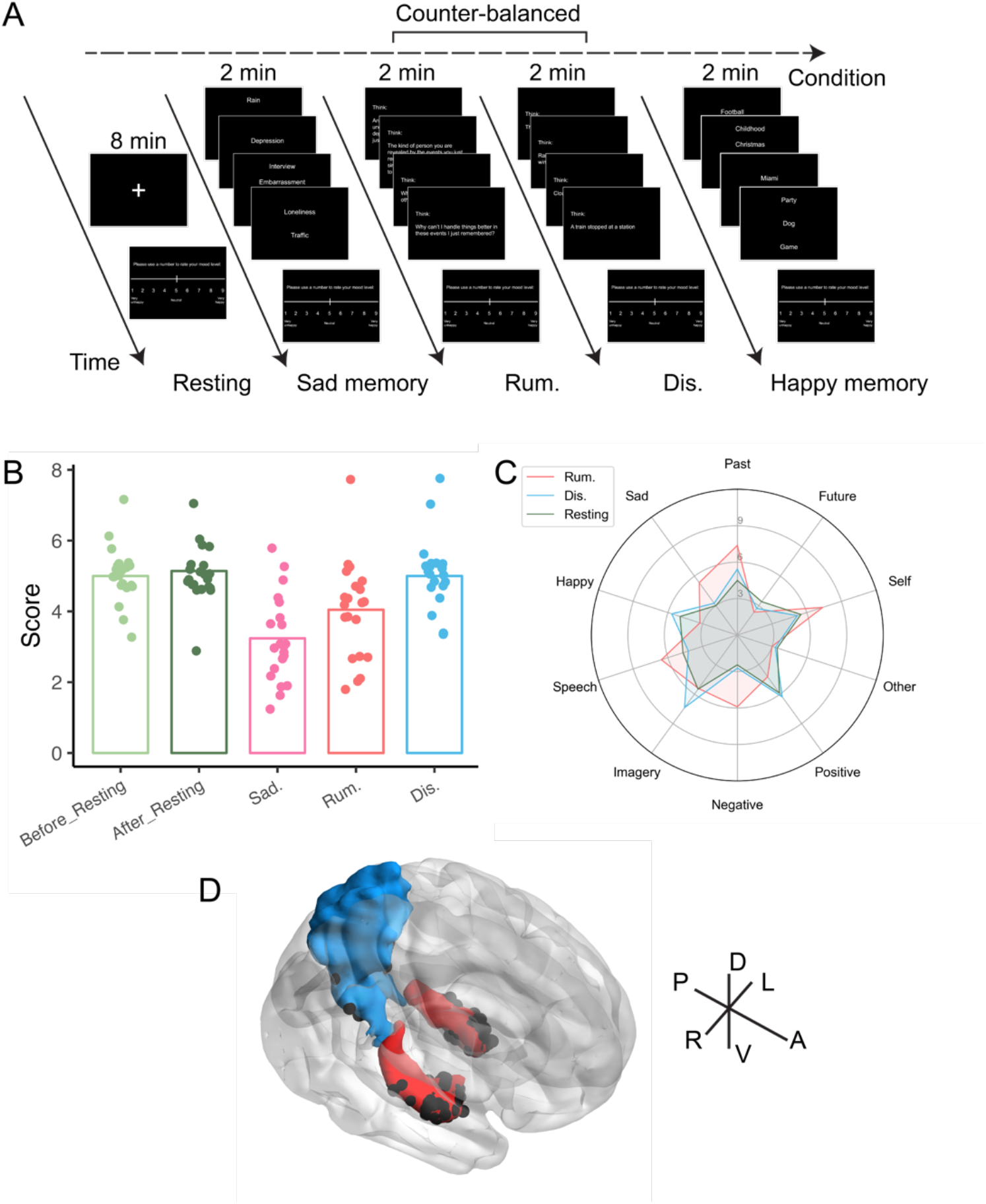
The rumination state task in the current intracranial electroencephalogram (iEEG) study. (A) The flow chart depicting the design of the rumination state task. (B) The emotional levels after each condition. (C) The form and content of thinking during rumination, distraction, and resting conditions. (D) All the depth electrodes located in the precuneus and the parahippocampal gyrus.

**Figure 2.**
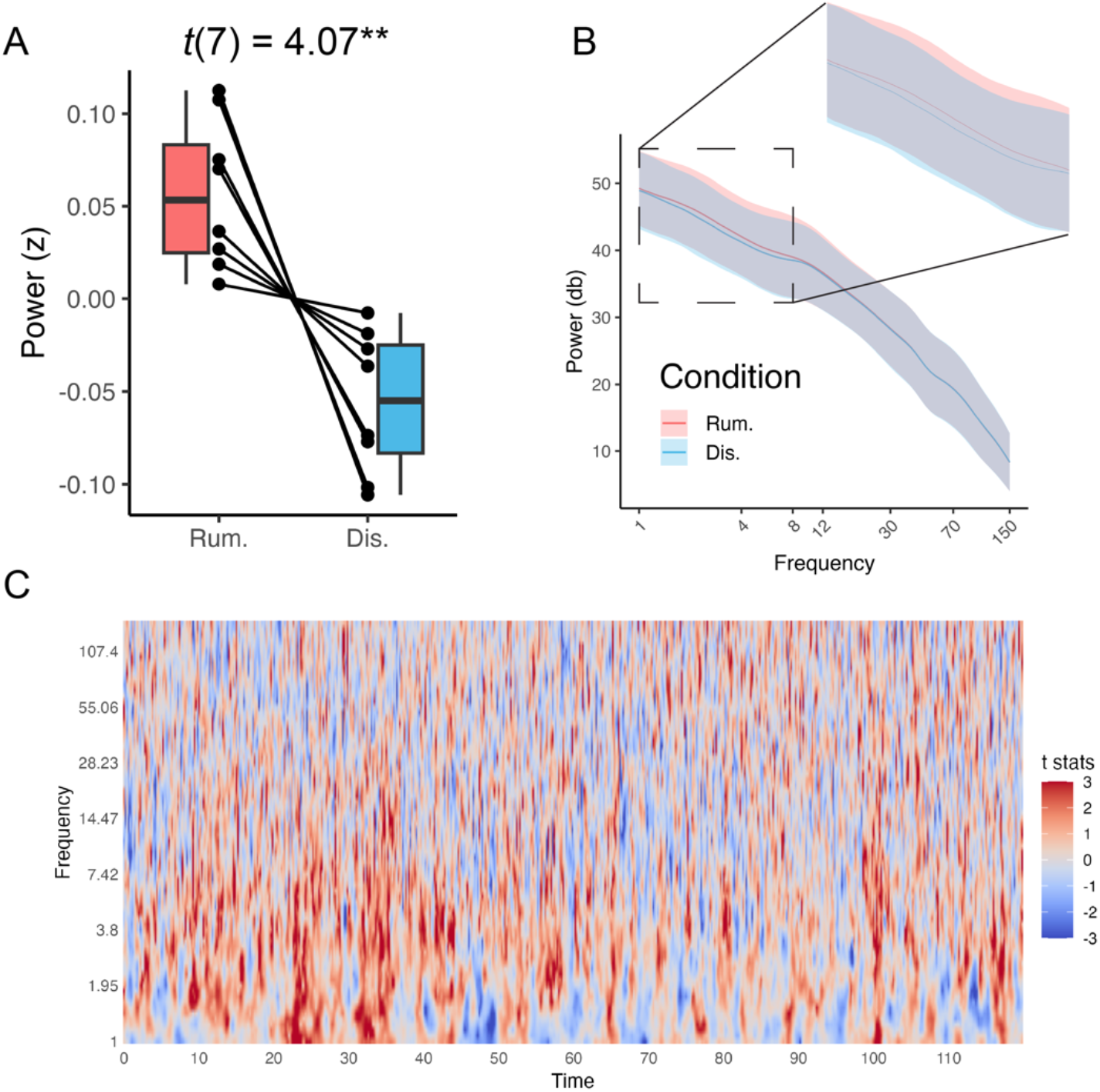
The intracranial electroencephalogram (iEEG) power changes in the precuneus. (A) Slow-frequency power during rumination was higher than in the distraction condition. (B) The Power spectrum is averaged across time during rumination and distraction conditions. The grey ribbon denotes the standard deviation of the average power across all participants. (C) The dynamic power changes across time during rumination compared to distraction. **: *p*_adj_ < .01

### Oscillation power during rumination as compared to distraction conditions

We observed a dissociated iEEG response pattern in the precuneus and hippocampus. In the rumination condition, the precuneus had an enhanced low-frequency band power compared to the distraction condition (*t*(7) = 4.07, *p*_*adj*_ = .01, ***η***^2^ = .70) (Figure 2A). The power spectra and the spectrogram of a typical block are shown in Figure 2. The average power throughout the entire time series in the distraction condition was used as the control to generate the spectrogram.

We observed a general and continuously enhanced slow-frequency power during an active rumination compared to distraction in the precuneus. A decreased high gamma band power was revealed in the hippocampus during rumination compared to distraction (*t*(18) = -2.77, *p*_*adj*_ = .02, ***η***^2^ = .30) (Figure 3A). Figure 3 shows the power spectra and the spectrogram of a typical block in the hippocampus. A weaker but similarly consistent reduction in power was observed in the hippocampus during rumination compared to distraction.

**Figure 3.**
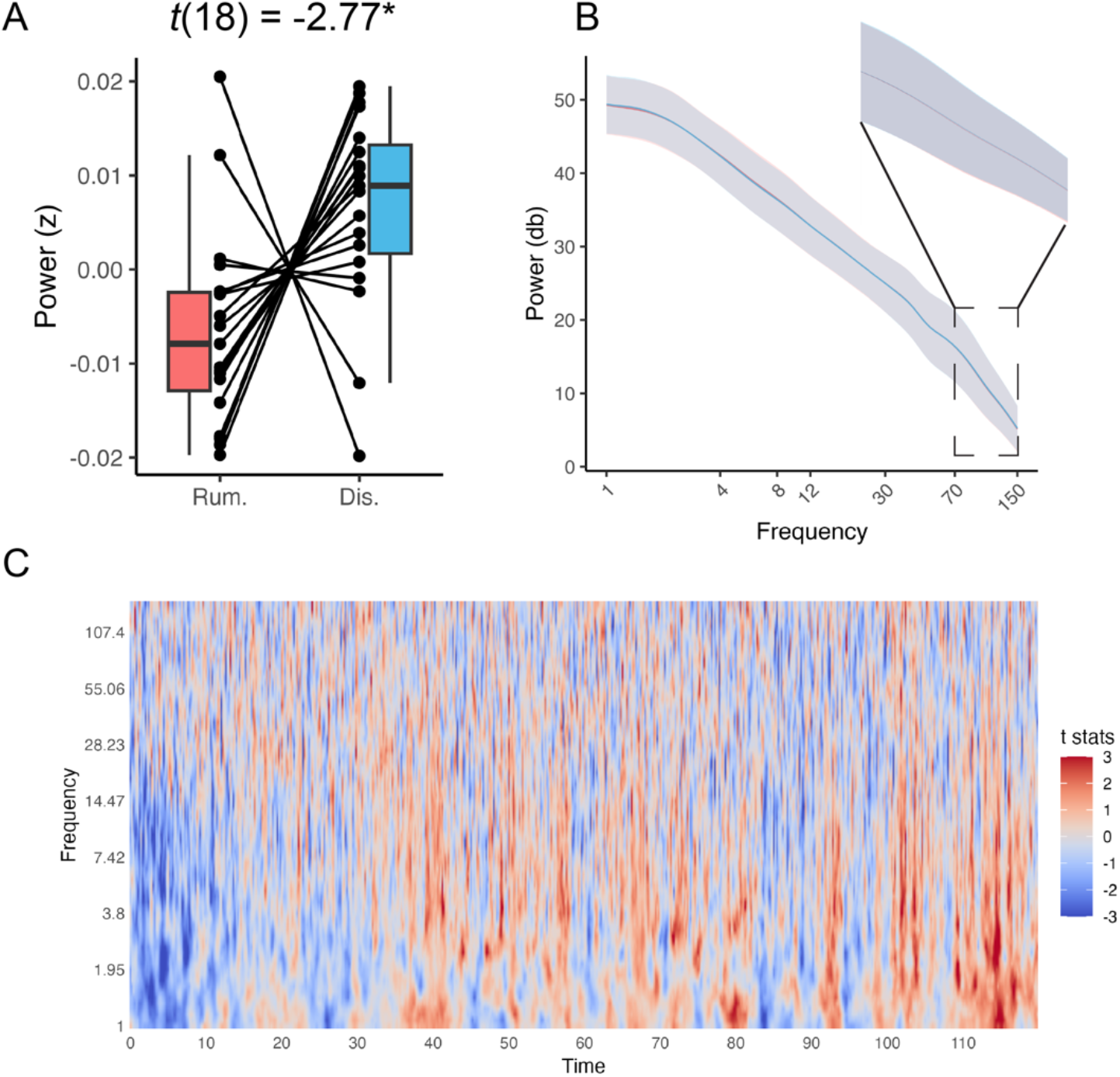
The intracranial electroencephalogram (iEEG) power changes in the hippocampus. (A) High gamma power during rumination was higher than in the distraction condition. (B) The Power spectrum is averaged across time during rumination and distraction conditions. The grey ribbon denotes the standard deviation of the average power across all participants. (C) The dynamic power changes across time during rumination compared to distraction. *: *p*_adj_ < .05

### Relationship between oscillation powers and self-reported rumination tendency

We observed a significant positive correlation between the delta high gamma power (rumination vs. distraction) in the hippocampus and the self-reported reflection tendency (*r*(13) = .71, *p*_*adj*_ = .01) (Figure 4).

**Figure 4.**
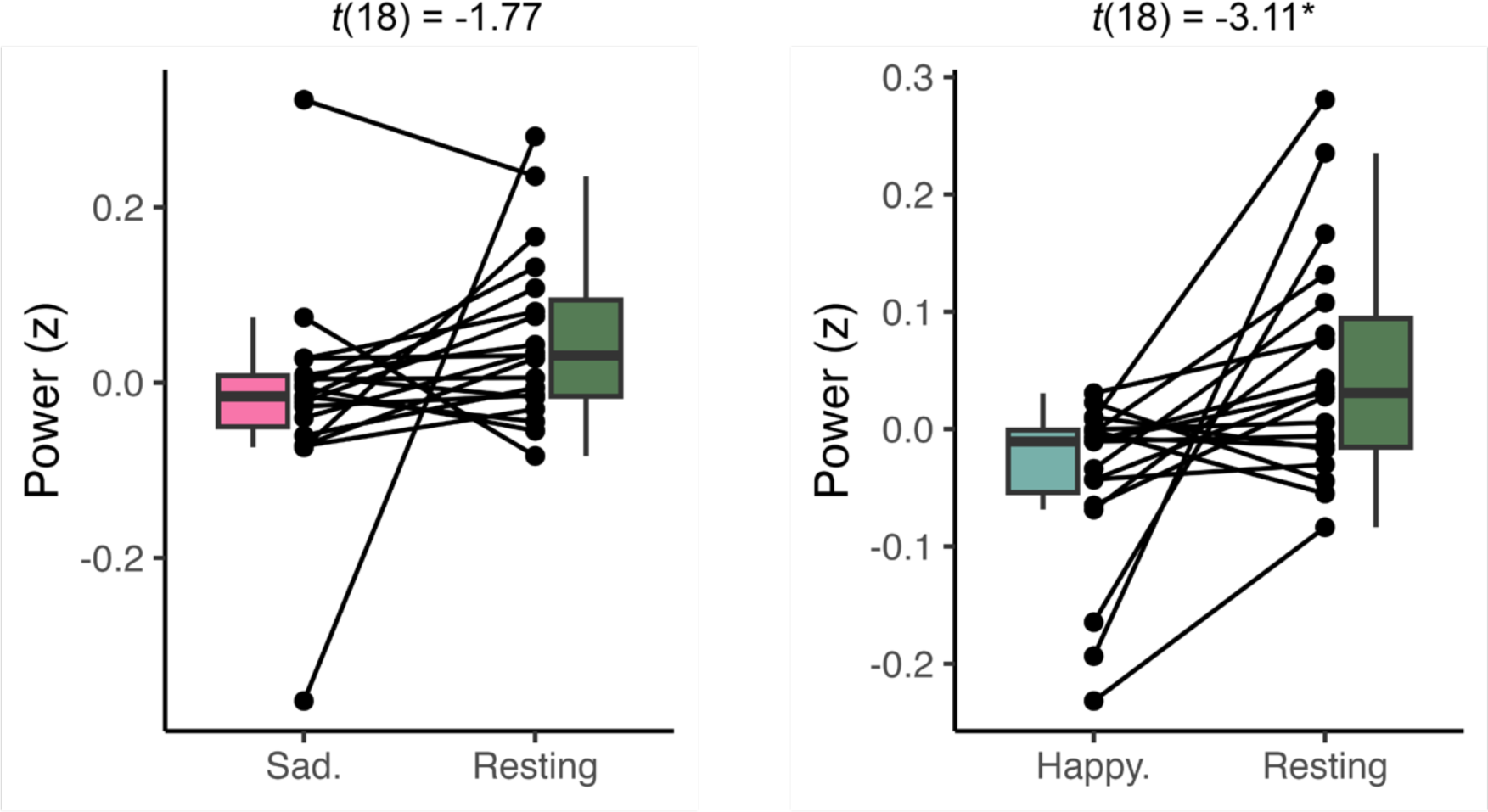
The hippocampus’s slow frequency band power significantly decreased during autobiographical memory conditions compared to the resting state. Abbreviations: Sad.: sad memory; Happy. Happy memory. *: *p*_adj_ < .05

**Figure 4.**
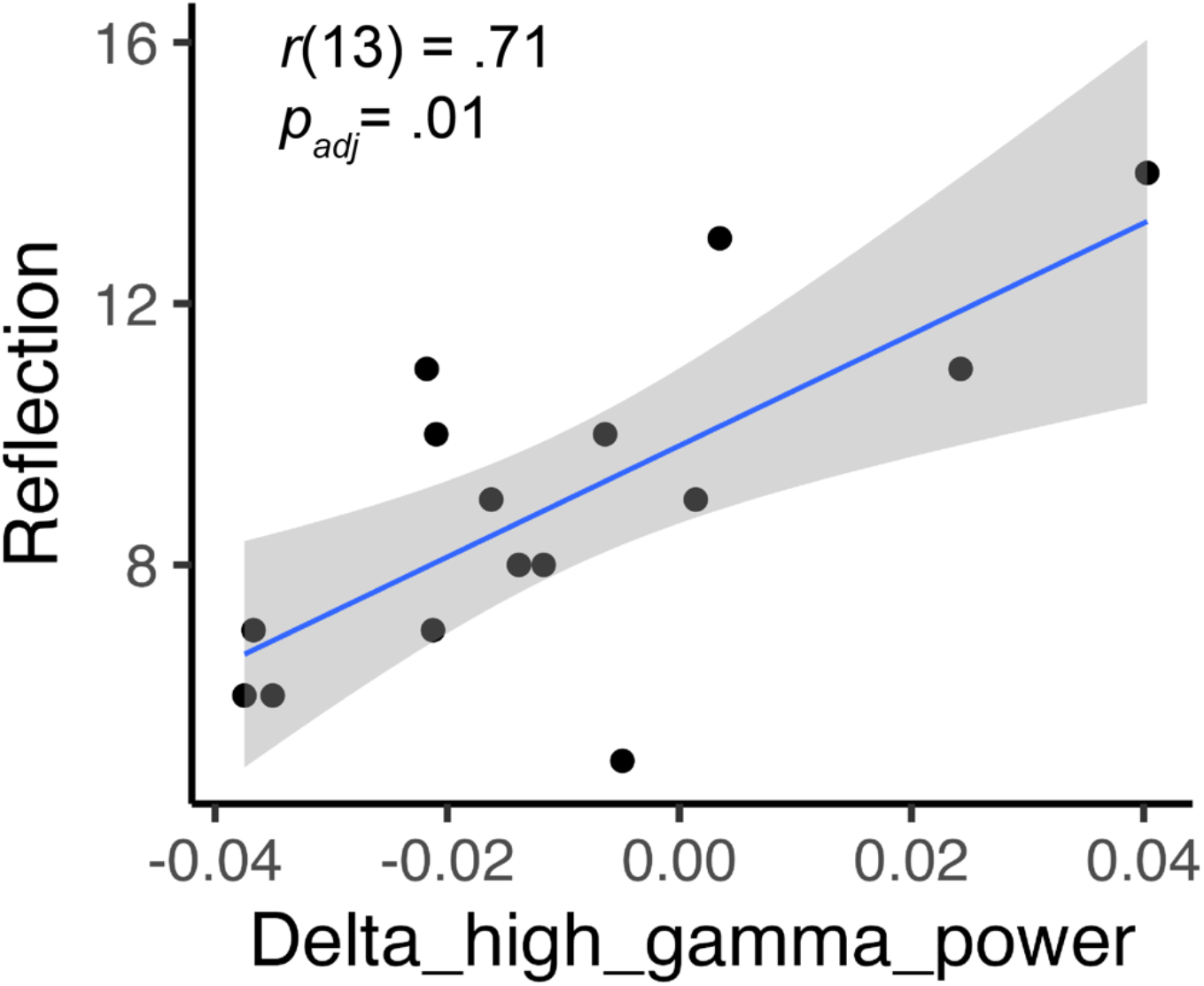
Relationship between the self-reported reflection tendency and the delta high gamma power (rumination vs. distraction) in the hippocampus. Note that four participants were excluded due to missing values of self-reported reflection scores.

Oscillation power during the autobiographical memory condition and resting state To verify whether participants have followed the prompts in the autobiographical memory conditions, we compared the averaged oscillation band powers in the hippocampus in autobiographical memory conditions compared to the resting state. We found a reduced slow-band frequency power in the hippocampus during the happy autobiographical memory condition compared to the resting condition (*t*(18) = -3.10, *p*_*adj*_ = .01, ***η***^2^ = .35). The same effect was revealed between the sad autobiographical memory condition but did not achieve significance (*t*(18) = -1.77, *p*_*adj*_ > .05, ***η***^2^ = .15)(Figure 4).

### Sensitivity analyses

We didn’t observe any significant difference in the slow frequency or high gamma powers between rumination and distraction conditions in the middle occipital gyrus (all *p*s > .05, Figure 5). The precuneus’ slow-frequency powers of all 4 prompts during the rumination condition were significantly higher than the distraction condition (all *p*s < .05, Figure S1). The high gamma powers of all prompts in the hippocampus decreased during rumination compared to the distraction, though the two prompts did not achieve significance.

**Figure 5.**
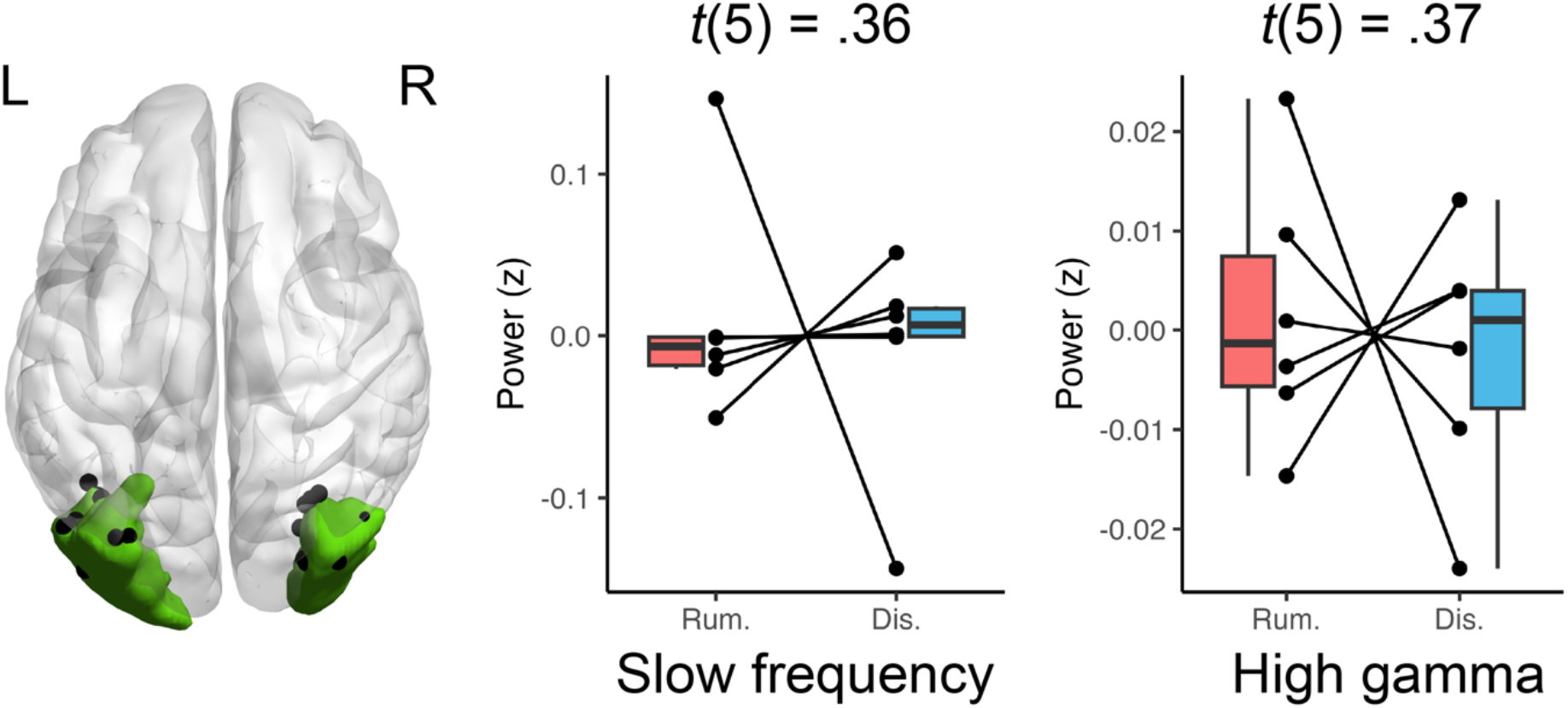
The slow-frequency/high gamma oscillation power change in the middle occipital gyrus during the rumination condition compared to the distraction condition.

## Discussion

This study provides the first empirical evidence of the direct electrophysiological basis underlying rumination in humans. We found dissociated power changes in the precuneus and hippocampus compared to the control condition during an active rumination state. Specifically, an enhanced slow frequency band power in the precuneus and a reduced high gamma band power in the hippocampus region during the rumination condition compared to the distraction condition. We further showed that the high-gamma power reduction in the hippocampus was associated with the tendency for self-reported reflection. Sensitivity analysis showed that the visual region (occipital lobe) was not involved in the neurophysiological processes underlying rumination.

Our results add to the existing fMRI studies, highlighting the prominent roles of the hippocampus and precuneus in the neural underpinnings of rumination (*8-10, 19, 21, 25*) and shed new light on the possible electrophysiological basis of such involvement. The enhanced slow frequency band power during rumination we found in the current study is in line with one previous scalp electroencephalogram study (*51*). In this study, Anderson and colleagues showed increased 4-6 Hz EEG power in the parietal regions during rumination, which could originate from the precuneus. Our results directly proved this notion using iEEG recordings. Slow frequency oscillations, especially the so-called theta band (4-8 Hz), have been well characterized to underlie spatial representation, memory, and consolidation in animal studies (*52*). Previous iEEG studies in humans also highlighted its involvement in the processes of learning and memory (*34, 35, 53-55*). Slow frequency oscillations may be a mechanism by which widespread neural assemblies communicate by synchronizing the precise timing of neuro spikings (*56, 57*). It should be noted that there is an ongoing discussion on rodent theta oscillation’s correspondence in humans, with some studies showing that a lower range of frequencies (i.e., 1-4 Hz) in humans was involved in the theta-band-related processes in rodents (*48, 58*). On the other hand, high gamma band oscillations have been suggested to be related to local neural population spikings and shown to be the possible neurophysiological basis for BOLD signals (*59, 60*). The hippocampus has been found to be critical to episodic and working memory in animal and lesion studies, as well as iEEG studies in humans (*61-65*). Rumination is closely related to negative biases in memory (*66*). Ruminative individuals tend to recall categorical memories lacking specific details, which has been concepted as overgeneral autobiographical memory (OGM) (*67-69*). Mctiormick and colleagues (*70*) examined the characteristics of the spontaneous thought of a group of human subjects with hippocampus lesions and found that they engaged in more verbally-sematic, abstract, and less imaginary, concrete thoughts compared to healthy controls. This rare, causal evidence implicates that the suppression of hippocampal activity may lead to ruminative thought that is repetitive, abstract, and in a verbal form. Aligning with this notion, we found a reduced high-gamma oscillation power in the hippocampus during the rumination condition.

Our results further implicated a possible interaction between the hippocampus and the precuneus during the rumination condition. The precuneus region, especially the posterior cingulate cortex, is a major target of the anatomical hippocampal efferents (*71*). Evidence from animal studies has shown that the precuneus-hippocampus coupling may be supported by the theta oscillations (*72*). Thus, a possible explanation of our observations is that the precuneus exerts cognitive control through theta oscillation and leads to a suppression of high gamma oscillation power in the hippocampus during an active rumination state. This implication is in line with the existing theoretical model of rumination, which argues that rumination is basically a tendency to process autobiographical information in a biased, abstract way, and this tendency may be due to a rigid constraint from neocortical regions such as precuneus to the medial temporal regions including the hippocampus (*1, 73, 74*). Apart from these models, which are based primarily on human studies, our findings also align with a recent hypothetical model based on animal studies, which states that the reverse hippocampal replay, a rapid, coordinated reactivation of encoding-activated neural ensembles, may be the neurophysiological basis for rumination (*75*). Nevertheless, the current study did not directly examine the possible coupling between these two regions, and future studies are needed to dwell on this intriguing direction.

We found a reduced slow-frequency band power in the hippocampus during the autobiographical memory condition relative to the resting state. The hippocampus has long been denoted to be closely involved in the episodic memory process (*76*). Evidence from lesion studies, including the famous case of H. M., has implicated the vital role of the hippocampus in encoding information and forming new episodic memories (*77, 78*). Specifically, the slow band power, especially the theta band power, has been found to be altered when rodents are performing memory tasks (*56, 79*). In line with the present study, most iEEG studies in humans found reduced slow frequency band (theta) oscillation power during episodic memory tasks (*80-82*). It is worth noting that, unlike most previous studies, which conducted typical working memory tasks and asked participants to remember a list of stimuli and try to retrieve them after a short period of time. We conducted a more naturalistic, self-constraint autobiographical memory task. Rudlor and colleagues (*83*) recently showed that this theta power reduction may be associated with broadband activity. The specific experimental design of the present study may allow for a more comprehensive and dynamic test of this notion. We observed a significant correlation between the degree to which the high-gamma band oscillation power decreased in the rumination condition compared to the distraction condition and the self-reported reflection tendency. Although deemed as a subtype of rumination, reflection is conceptualized as a purposeful self-pondering to engage in problem-solving to alleviate an individual’s dysphoric symptoms (*84, 85*). Unlike other subtypes, brooding and reflection might be more constructive in terms of mental health. Accordingly, the reduced hippocampal high gamma high-gamma band activity may lead to more abstract, less constructive thinking and make one’s self-pondering less like reflection.

One of the strengths of the present study is the consistency across subjects. Leveraging the extensive electrode coverage of the two ROIs from a relatively large sample, we demonstrated an ROI-wised, subject-level effect instead of electrode-level findings. We also applied a more naturalistic, continuous, non-constrained experimental design, which is more suitable for rumination. As one abnormal form of spontaneous thought, rumination exhibits variable temporal dynamics and may not be time-locked to any explicit external events (*20*). The validation analysis in a typical visual region further showed the specificity of our findings to the precuneus and hippocampal local neurophysiological activities.

Our findings should be interpreted with respect to limitations. First, due to the explorative nature of the present research, we only examined the averaged local oscillation powers, omiting the possible coupling between the precuneus and the hippocampus. Existing evidence has shown that the phase of low-frequency oscillations is co-modulated with high gamma activity during both resting and task states (phase-amplitude coupling, PAC) (*86-89*). Hebscher and colleagues examined the PAC between the precuneus and the medial temporal lobe during an autobiographical memory task and found its association with memory retrieval (*90*). More importantly, they later successfully changed the vividness of the memory by modulating such PAC with transcranial magnetic stimulation (TMS) to the precuneus. In light of a recent study showing the effects of theta burst stimulation (TBS), a novel TMS protocol that functions by mimicking the brain’s theta band activity (*91*), can be enhanced by adapting to an individual’s PAC pattern (*92*). Future studies could develop a targeted TMS protocol for the precuneus/hippocampus oscillation activities and tailor the parameters according to the individual’s specific PAC pattern between these two brain regions. Second, the results reported here were based on a group of patients with epilepsy who showed moderate depressive symptoms, so it is cautious to generalize our findings to other clinical samples, such as patients with major depressive disorder (MDD). Our previous function MRI research has revealed a case-control difference regarding the rumination’s brain network mechanism (*24, 93*), albeit the effect size was moderate, and a noticeable similarity was also found.

In conclusion, using iEEG recordings in a group of patients with epilepsy, we found an enhanced slow frequency band power in the precuneus and a decreased high gamma power in the hippocampus in a naturalistic, continuous, active rumination state. Further analysis showed its relevance to self-reported reflection and specificity to these two brain regions. Our findings provided the first empirical evidence of the underlying neurophysiological mechanism of rumination and implicated a possible control from the precuneus to the hippocampus through neural oscillation coupling during an active rumination.

## Acknowledgments

Xiao Chen has received research support from the National Natural Science Foundation of China (No. 32300933), and the China Scholarship Council (CSC, No. 202104910248).

