## Supplementary_materials for "Intracranial neurophysiological correlates of rumination"

Table S1. The prompts to induce rumination and distraction.

| **Prompts** | **Condition** |
| --- | --- |
| Think: Analyze your personality to understand why you feel so depressed in the events you just remembered | Rumination |
| Think: The kind of person you are revealed by the events you just remembered. How similar/different you are relative to other people？ | Rumination |
| Think: Why do I encounter these events other people don’t？ | Rumination |
| Think: Why can’t I handle things better in these events I just remembered？ | Rumination |
| Think about: The layout of a typical classroom | Distraction |
| Think about: Raindrops sliding down a windowpane | Distraction |
| Think about: Clouds forming in the sky | Distraction |
| Think about: A train stopped at a station | Distractions |

Table S2. The self-reported thinking forms and contents during the resting state and the rumination/distraction conditions

| **Characteristics of thinking contents** | **resting** | **rumination** | **distraction** | ***F*** | ***p*** |
| --- | --- | --- | --- | --- | --- |
| Past | 4.50 (2.04) | 7.38 (1.72) | 5.43 (2.66) | 7.85^ab^ | .001 |
| Future | 3.40 (2.06) | 2.33 (1.49) | 2.71 (2.19) | 1.83 | .17 |
| Myself | 5.50 (2.44) | 7.38 (2.18) | 5.19 (2.25) | 6.76^a^ | .01 |
| Others | 3.45 (2.04) | 3.00 (1.84) | 3.29 (2.03) | .30 | .74 |
| Positive | 5.90 (2.22) | 4.24 (2.53) | 6.29 (1.76) | 6.49^a^ | .004 |
| Negative | 2.45 (2.06) | 5.90 (2.19) | 2.71 (1.62) | 23.81^ab^ | < .001 |
| Image | 5.50 (2.98) | 5.43 (2.66) | 7.38 (2.01) | 6.84^ac^ | .003 |
| Speech | 4.70 (3.26) | 6.57 (2.79) | 4.24 (2.57) | 6.64^a^ | .003 |
| Happy | 4.95 (1.85) | 3.24 (2.12) | 5.67 (1.83) | 8.49^ab^ | < .001 |
| Sad | 3.00 (1.97) | 5.33 (2.15) | 3.24 (1.70) | 13.14^ab^ | < .001 |
| All values are mean (SD) | | | |  |  |

a: significant differences between the rumination and distraction conditions

b: significant differences between the rumination and resting conditions

c: significant differences between the distraction and resting conditions

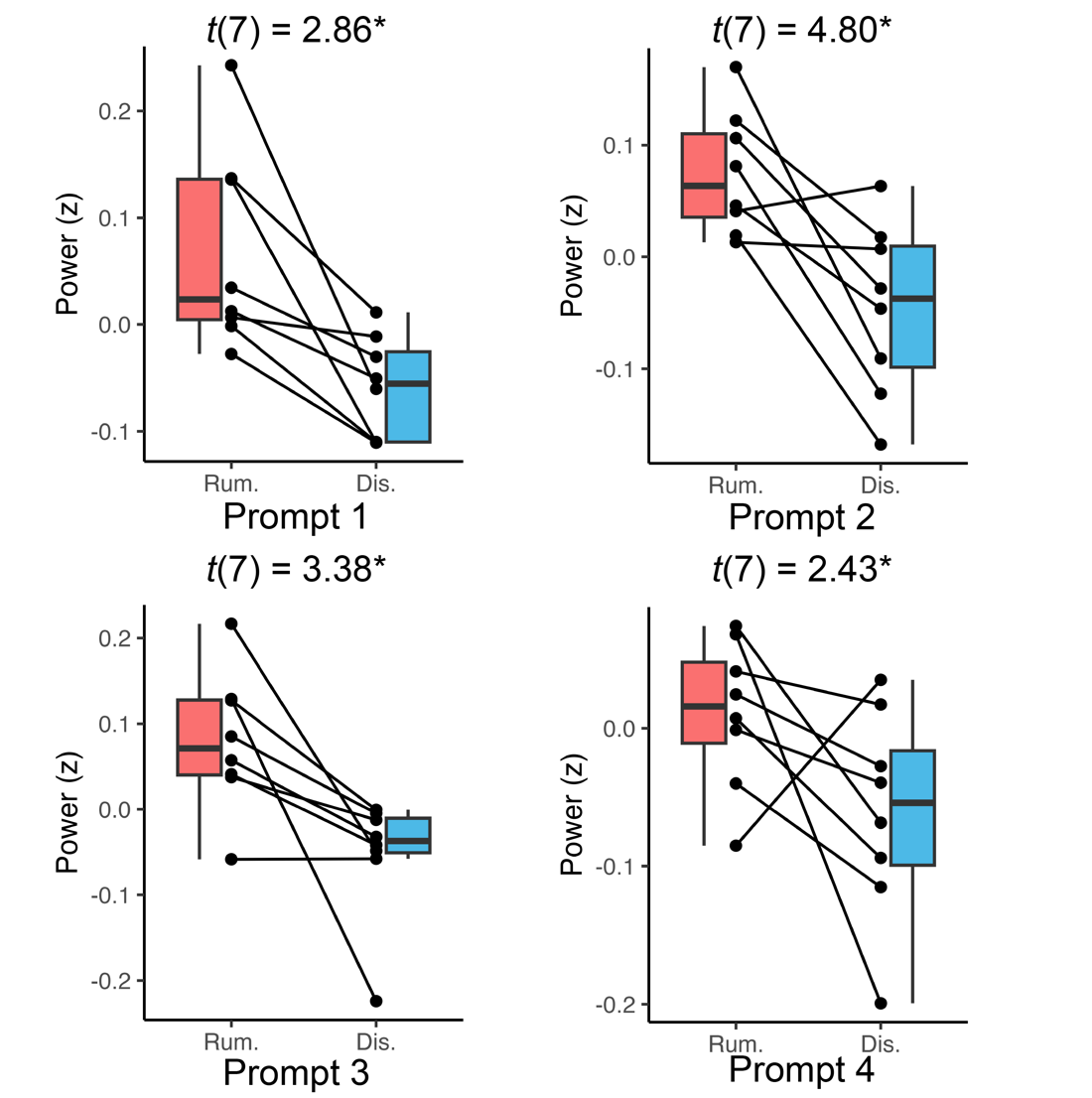

Figure S1. Slow-frequency power in the hippocampus during the rumination condition compared to the distraction condition.

Abbreviations: rum., rumination; dis., distraction.

*: *p* < .05

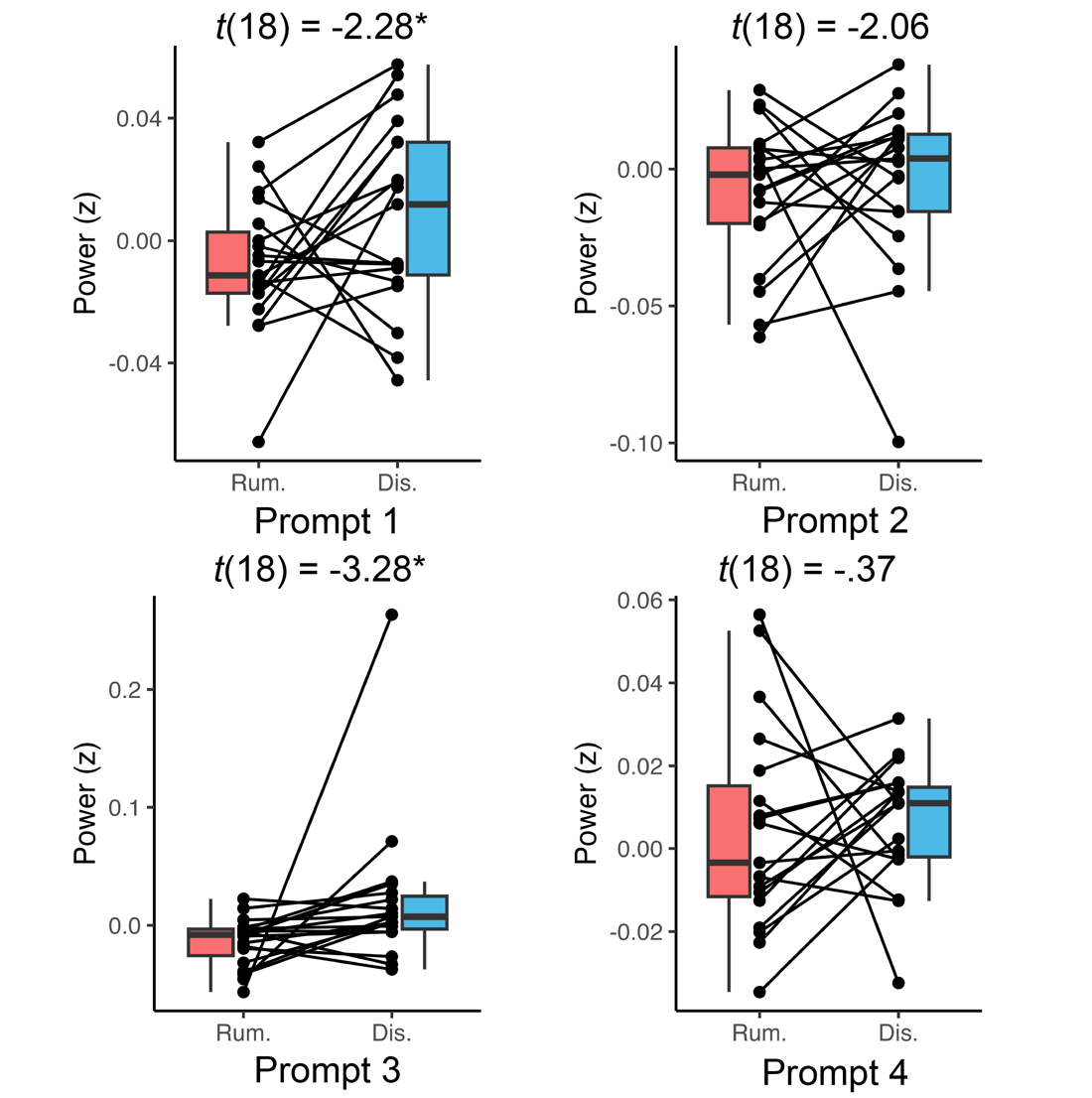

Figure S2. High gamma power in the hippocampus during the rumination condition compared to the distraction condition.

Abbreviations: rum., rumination; dis., distraction.

*: *p* < .05
